# ATP Citrate Lyase Supports Cardiac Function and NAD+/NADH Balance And Is Depressed in Human Heart Failure

**DOI:** 10.1101/2024.06.09.598152

**Authors:** Mariam Meddeb, Navid Koleini, Seungho Jun, Mohammad Keykhaei, Farnaz Farshidfar, Liang Zhao, Seoyoung Kwon, Brian Lin, Gizem Keceli, Nazareno Paolocci, Virginia Hahn, Kavita Sharma, Erika L. Pearce, David A. Kass

**Affiliations:** Division of Cardiology, Department of Medicine, Johns Hopkins University School of Medicine, Baltimore, MD; Department of Department of Oncology, Department of Biochemistry and Molecular Biology, Johns Hopkins University School of Medicine, Baltimore, MD; Department of Pharmacology and Molecular Sciences, Johns Hopkins University, Baltimore MD

## Abstract

**Background:** ATP-citrate lyase (ACLY) converts citrate into acetyl-CoA and oxaloacetate in the cytosol. It plays a prominent role in lipogenesis and fat accumulation coupled to excess glucose, and its inhibition is approved for treating hyperlipidemia. In RNAseq analysis of human failing myocardium, we found *ACLY* gene expression is reduced; however the impact this might have on cardiac function and/or metabolism has not been previously studied. As new ACLY inhibitors are in development for cancer and other disorders, such understanding has added importance.

**Methods:** Cardiomyocytes, *ex-vivo* beating hearts, and *in vivo* hearts with ACLY inhibited by selective pharmacologic (BMS303141, ACLYi) or genetic suppression, were studied. Regulation of ACLY gene/protein expression, and effects of ACLYi on function, cytotoxicity, tricarboxylic acid (TCA)-cycle metabolism, and redox and NAD+/NADH balance were assessed. Mice with cardiac ACLY knockdown induced by AAV9-*acly-*shRNA or cardiomyocyte tamoxifen-inducible *Acly* knockdown were studied.

**Results:** *Acly* gene expression was reduced more in obese patients with heart failure and preserved EF (HFpEF) than HF with reduced EF. *In vivo* pressure-overload and *in vitro* hormonal stress increased ACLY protein expression, whereas it declined upon fatty-acid exposure. Acute ACLYi (1-hr) dose-dependently induced cytotoxicity in adult and neonatal cardiomyocytes, and caused substantial reduction of systolic and diastolic function in myocytes and *ex-vivo* beating hearts. In the latter, ATP/ADP ratio also fell and lactate increased. U_13_C-glucose tracing revealed an ACLY-dependent TCA-bypass circuit in myocytes, where citrate generated in mitochondria is transported to the cytosol, metabolized by ACLY and then converted to malate to re-enter mitochondria, bypassing several NADH-generating steps. ACLYi lowered NAD+/NADH ratio and restoring this balance ameliorated cardiomyocyte toxicity. Oxidative stress was undetected with ACLYi. Adult hearts following 8-weeks of reduced cardiac and/or cardiomyocyte ACLY downregulation exhibited ventricular dilation and reduced function that was prevented by NAD augmentation. Cardiac dysfunction from ACLY knockdown was worse in hearts subjected to sustained pressure-overload, supporting a role in stress responses.

**Conclusions:** ACLY supports normal cardiac function through maintenance of the NAD+/NADH balance and is upregulated by hemodynamic and hormonal stress, but depressed by lipid excess. ACLY levels are most reduced in human HFpEF with obesity potentially worsening cardio-metabolic reserve.

## Introduction

In 2023, The World Health Organization reported nearly 16% of the world’s population lives with obesity^1^, the number being ≍40% in the United States^2^, and this pandemic is paralleled by an increased prevalence of metabolic syndrome (MS)^3, 4^. The consequences on cardiovascular health are substantial, as obesity is linked to hypertension, hyperlipidemia, atherosclerosis, and heart failure^5^. The association of obesity and heart failure is the strongest even after adjusting for other co-morbidities^6^, and their combination is often present in patients with heart failure and preserved ejection fraction (HFpEF)^7, 8^. Among factors linking obesity/MS to heart failure are dyslipidemia, lipid homeostasis, and altered metabolism, often coupled to metabolic dysfunction-associated steatotic-liver disease (MASLD)^9–11^. Obesity, MASLD and dyslipidemias are now targets for HF therapy^12–14^.

One enzyme that is gaining interest is ATP-citrate lyase (ACLY), a key controller of *de novo* lipogenesis, linking glucose to lipid metabolism^15^. ACLY is a highly conserved homo-tetrameric enzyme found in both the cytosol and the nucleus that converts citrate and co-enzyme A (CoA) into acetyl-CoA and oxaloacetate (OA)^16^. Citrate is generated in mitochondria as part of the tricarboxylic acid cycle (TCA) but it can also be transported to the cytosol and metabolized by ACLY^17^. ACLY-generated acetyl-CoA is used for fatty acid and cholesterol biosynthesis and as a substrate for protein acetylases, including histone acetyltransferases in the nucleus ^18^. ACLY generated OA can be converted to malate, which then re-enters mitochondria short-circuiting the canonical TCA cycle^19^. This alternative pathway bypasses two intermediate TCA steps that normally generate reduced nicotinamide adenine dinucleotide (NADH) required for oxidative phosphorylation. Cancer cells often upregulate ACLY, depleting cytosolic citrate and NADH to thereby augment glycolysis even in the presence of oxygen (Warburg effect), and provide glycolytic intermediates used for biosynthesis and acetyl-CoA for protein acetylation and membrane biosynthesis^20–22^. Non-proliferating differentiated cells are thought to mostly rely on the canonical TCA cycle^19^.

While ACLY is broadly expressed, it is most prominent in hepatic and adipose cells that store and process lipids. Homozygous *Acly* deletion is embryonically lethal, while mice with heterozygous deletion are viable and have little basal phenotype ^23^. Targeted deletion in adipose cells induces their resistance to insulin ^24^, whereas deletion in hepatocytes reduces NAFLD and associated dyslipidemia^25^. Mendelian randomization studies found human ACLY loss-of-function mutations associated with reduced risk of hyperlipidemia, atherosclerosis, and coronary disease ^26^. These findings spawned human use of bempedoic acid, an ACLY-inhibiting prodrug that requires activation by the liver, to lower low-density lipoprotein ^27^. Bempedoic acid is also reported to reduce MASLD in a preclinical model^25^, and new ACLY inhibitors are being developed for cancer and metabolic diseases ^28^. As these agents evolve, it is important to consider potential effects of ACLY inhibition on non-proliferative, oxidative, and catabolic tissues, such as the heart.

There is remarkably little reported about the ACLY in the heart, but our interest was kindled after finding *ACLY* gene expression to be significantly reduced in human myocardium from patients with HF and a reduced or preserved ejection fraction (HFrEF, HFpEF) ^29^. This led to the question of what ACLY’s impact on cardiac function and metabolism was, in particular what were the consequences of its downregulation? Here, we show that acute and sustained ACLY inhibition depresses myocyte and intact heart function coupled to reduced NAD+/NADH ratio, while supplementing myocytes or hearts with NAD+ corrects functional defects and redox balance.

## Methods

### Neonatal rat cardiomyocytes

Neonatal rat cardiomyocytes (NRCM) were isolated as previously described^30^. To disrupt ACLY, we used either a selective pharmacologic inhibitor (ACLYi) BMS303141 (Tocris #4609), reconstituted in DMSO at concentrations specified for each experiment, or silencing RNA gene knock-down. For studies using siRNA to downregulate *Acly*, myocytes were transfected in serum-free, antibiotic-free DMEM medium supplemented with ITS, with 100 nM of either ON-TARGETplus Non-targeting Control siRNA Pool (Dharmacon #D-001810-10-05) or On-TARGETplus Smartpool siRNA targeting rat ACLY protein (Dharmacon #L-089773-02-0005) using lipofectamine RNAimax (Thermofisher Scientific #13778075), as per manufacturer’s protocol.

For studies in which myocytes were exposed to lipids, chemically defined lipid concentrate (Thermofisher Scientific #11905031, Waltham, MA) or an equivalent volume of normal saline (vehicle) was added to the culture medium at a ratio of 1:100 of the total volume of medium. Endothelin-1 (100 nM, 24 hrs Sigma-Aldrich #E7764) was used to stimulate hypertrophy. Studies with added α-ketoglutarate were performed with addition of 2.5 mM of dimethyl-α-ketoglutarate (DMKG; Cayman chemical #28394) pre-buffered with sodium bicarbonate and HEPES (GIBCO #15630130) to target PH of 7.4 +/-0.2, or equivalent volume of normal saline with HEPES (vehicle). For studies done with N-acetylcysteine, 200 μM of N-acetyl-L-cysteine (Sigma Aldrich #A9165) was added to the culture medium for 4 hours prior to the initiation of treatment with ACLYi at 25 μM dose.

### Adult cardiac cell isolation

Adult mouse cardiomyocytes (AMCM) were isolated by retrograde intra-left ventricular perfusion of collagenase solution with aortic clamping as described^31^. Myocytes were plated in M199 media containing 10% FBS for 1 hour. Then media was changed to serum-free M199 supplemented with ITS and 2,3-butandione monoxime (BDM; Sigma-Aldrich #B0753).

### Metabolic studies

Isotope tracing was performed in freshly isolated AMCM plated in 6 well plates. After 4 hours of stabilization, cells were washed with PBS and changed into a glutamine and glucose-free M199 medium supplemented with 5 mM of [U-^13^C] glucose, 0.68 mM of glutamine, 1:100 CD lipids, with the addition of 10 μM of ACLYi or equivalent volume of DMSO (vehicle) for 1 hour prior to harvest. Untargeted metabolomics was performed from whole mouse heart (Langendorff preparation or explanted immediately following euthanasia) that were rinsed in PBS and immediately flash-frozen in liquid nitrogen. Metabolites were extracted in pre-cooled (-80°C) 80% HPLC grade methanol (Sigma-Aldrich #C34860). Metabolomic assays were performed either at the University of Pennsylvania Metabolomics Core, or in our laboratory (ELP).

ATP/ADP ratio was assessed with EnzyLightTM ADP/ATP Ratio Assay kit (BioAssay Systems #ELDT100), and NAD+/NADH ratio by NAD/NADH-Glo^TM^ kit (Promega #G9072) following manufacturer’s instructions.

### Assessment of NAD+/NADH ratio

For the assessment of mitochondrial NAD+/NADH balance, NRCM were transduced with a mitochondrial HA tag^32^ packaged into an adenovirus vector. Forty-eight hours later, cells were treated with 25 μM of ACLYi or equivalent volume of vehicle for 1 hour. One hour after treatment, cells were washed with PBS, lifted in fresh PBS with a cell lifter (Corning #3008) and mitochondria were isolated using Pierce^TM^ anti-HA magnetic beads (Thermo Scientific #88836) according to the manufacturer’s recommendation in the cold room. Isolated mitochondria were subsequently lysed and NAD+/NADH ratio immediately measured using the NAD/NADH-Glo^TM^ assay as described above.

### Cell viability and cytotoxicity assays

Cell viability in AMCM was assessed using the LIVE/DEAD Viability/Cytotoxicity Kit for mammalian cells (Thermofisher Scientific #L3224), and in both AMCM and NRCM by LDH release to the media with LDH-glo^TM^ assay (Promega) per manufacturer’s instructions.

### *Ex vivo* and *in vitro* functional studies

Rapidly excised mouse hearts (C57BL/6J) were mounted on a Langendorff preparation, with a balloon inserted into the left ventricle and constant volume set to generate a developed pressure of ∼90 mmHg. Details of the preparation have been previously described ^33^. Both male and female mice were used, aged 11±2 weeks, and they were randomly assigned to either 1μM of ACLYi ±2.5 mM DMKG, or to vehicle (DMSO). Rate pressure product (heart rate x developed pressure), and peak and trough of the first derivative of pressure were assessed continuously for 1 hour.

For myocyte function studies, freshly-isolated adult mouse cardiomyocytes were treated with 1μM of ACLYi or equivalent volume of DMSO. Immediately following 1 hour of treatment, cardiomyocyte function was evaluated using the IonOptix system to assess both sarcomere shortening and whole cell calcium transients, as previously described^34^.

### Mitochondrial respiration study

This was performed in NRCMs, with cells previously incubated with either scrambled or siRNA to ACLY (see main Methods). One hour prior to the metabolic flux measurement, the culture medium was substituted by DMEM Seahorse XF DMEM assay medium with 5 mM of glucose, 1 mM of pyruvate and 2 mM of glutamine, and cells were incubated at 37°C in a non-CO2 incubator for 1 hour prior to data acquisition. Oxygen consumption rate was measured at baseline and after serial injections of 1.5mM of oligomycin A (Sigma-Aldrich #75351), 2 mM of carbonyl cyanide-p-trifluoromethoxyphenylhydrazone (FCCP, Sigma-Aldrich #SML2959) and 2 mM of rotenone (Millipore Sigma #R8875) and antimycin (Sigma-Aldrich #A8674).

### Reactive Oxygen Species Analysis

#### Electron Paramagnetic Resonance (EPR)

NRCM were treated with 25 μM of ACLYi or vehicle for 1 hour. After incubation, ROS was analyzed by electron paramagnetic resonance (EPR) spectroscopy, as previously described ^35^. The EPR signal intensities were normalized to the protein concentrations of the samples as determined by BCA protein assay kit (Thermofisher Scientific^TM^).

#### Mitochondrial ROS

NRCM were treated with 25 μM of ACLYi or vehicle for 1 hour. Cells treated with 10 mM hydrogen peroxide for 5 minutes were used as a positive control. The culture medium was subsequently replaced with PBS including 100 nM of MitoSOX^TM^ red reagent (Invitrogen #M36008). After 15 minutes of incubation with the MitoSOX^TM^ reagent, cells were imaged with a Revolve microscope (ECHO) using the rhodamine filter. The intensity of mitochondrial fluorescence was then measured using ImageJ software. The calculated fluorescence intensity after subtraction of background fluorescence for individual cells was used to compare mitochondrial ROS between groups.

#### Cytosolic ROS

NRCM were transduced with cytoORP1; an adenovirus construct expressing redox-sensitive GFP (roGFP) fused with the yeast peroxidase ORP1, localizing to the cytosol in live cells, as previously described ^36^, courtesy of Dr Brian O’Rourke’s laboratory. After 48 hours of transduction, at which time point a transduction efficiency of >90% was achieved, cells were treated with 25 μM of ACLYi or equal volumetric ratio of DMSO (vehicle) for 1 hour. The medium was subsequently changed to a modified Tyrode solution (10 mM glucose, 2 mM pyruvate, 130 mM NaCl, 5 mM KCl, 1 mmol MgCl_2_, 10 mM HEPES, 2 mM CaCl_2_ and 0.3mM ascorbic acid) and cells were imaged with an Andor XD Revolution spinning disk confocal microscope (Olympus) using 405- and 488-nm laser lines for excitation and 500-to 554nm bandpass filter for detection. Images were acquired every 2 seconds over 30 minutes, after which signals were calibrated using diamide to obtain the R_max_ for roGFP probe and dithiothreitol (DTT) to obtain the R_min_. ROS emission was then estimated by the 405/488 nm excitation ratio images that were created and analyzed using ImageJ software after normalization of the ratiometric signals to the R_min_ and R_max_.

### Analysis of mRNA

RNA was extracted from cells and heart samples using TRIzol^TM^ Reagent (ThermoFisher Scientific #15596026), eluted in RNAse-free water and reverse transcribed with High Capacity cDNA Reverse Transcription Kit with RNase Inhibitor (Applied Biosystems #4374966) according to the manufacturers’ instructions. Semi-quantitative reverse transcription PCR was done on a QuantStudio^TM^ 5 Real-Time PCR (Thermofisher Scientific^TM^) with specific Taqman probes targeting the genes of interest using the TaqMan Fast Advanced Master Mix for qPCR (Applied Biosystems #4444963). Expression levels of the transcripts were normalized to the level of the housekeeping gene ribosomal protein S18 (RPS18) using the difference in Ct values between the gene of interest and RPS18. The relative changes in gene expression were compared between experimental and control conditions using the 2 ^-ΔΔCt^ method.

### Analysis of protein expression

Cells were scraped in SDS lysis buffer supplemented with 1:100 Halt^TM^ Protease and Phosphatase inhibitor cocktail (100X) (Thermo Scientific #78446). The lysates were sonicated, heated, and centrifuged (14,000g for 15 minutes) to remove debris. Protein concentration was quantified in the supernatants using the bicinchoninic acid assay (BCA). Following SDS-PAGE, proteins were transferred to nitrocellulose membranes. Membranes were blocked in Intercept® (TBS) Blocking Buffer (Li-COR) for 1 hour at room temperature to stop nonspecific binding. Membranes were incubated overnight at 4°C with the primary antibody diluted 1:1000 in Intercept® T20 (TBS) Antibody Diluent (Li-COR), washed in TBS-Tween (TBS-T) the following morning, followed by 1 hour incubation in secondary antibody at 1:5000 dilution at room temperature, followed by 1 hour of thorough washing in TBS-T and visualized by infrated imaging using Odyssey (Li-COR). Total protein concentration was determined using Revert^TM^ 520 Total Protein Stain kit (Li-COR) according to the manufacturer’s instructions.

### Generation of adenovirus for ACLY overexpression and promotor activity assay

Replication deficient adenovirus plasmids were linearized using PacI enzyme (NEB, Cat# R0547) as per manufacturer protocol. Human embryonic kidney 293T cells (ATCC, Cat # CRL-3216) were transfected by the linearized plasmids (10 mg per over 95% confluence T25 flasks) using Lipofectamine 3000 (Thermofisher, Cat# L3000001) as per manufacturer protocol. The next day, the media was renewed on the cells, and cells were cultured for 7 days. To acquire the virus, cells were collected and resuspended in 1 ml of PBS and subjected to 3 times of freeze/thaw cycles. Lysates were centrifuged at 2000g for 10 minutes, and supernatants were collected. The supernatants were put on >95% confluence HEK293T cells in a T75 flask and cultured for 7 days. Cells were collected, and an active virus was acquired. The virus titer was estimated by serial dilution as previously described ^37^. Cells were transduced with the virus at a multiplicity of infection (MOI) of 20.

### Promoter activity studies

NRVM were transduced with an adenovirus vector carrying the human ACLY promoter coupled to luciferase (Luc2) as a reporter gene and cultured at 37°C in DMEM/ITS medium with or without 1:100 Chemically Defined Lipid Concentrate (Gibco #11905031). Twenty-four hours later, cells were rinsed with PBS and lysed with 1X Passive Lysis Buffer (Promega #E1941). The luciferase assay reagent (Promega #E1501) was subsequently added to the cell lysates and the luciferase activity was measured using the Glomax 96 plate reader (BioTek), according to the manufacturer’s protocol.

### Assessment of mitochondrial membrane potential

NRCM were treated with 25mM ACLYi or vehicle for 1 hr. Cells were then stained with 20 nM TMRM staining solution, incubated for an additional 30 minutes and the mitochondrial membrane potential was subsequently assessed using the Tetramethylrhodamine, methylester, Perchlorate (TMRM) reagent (Invitrogen #T668), according to the manufacturer’s protocol. Fluorescence intensity of cells was quantified using image J (Fiji) and normalized to background.

### Fluorodeoxyglucose position electron tomography

Mice (age 12 weeks) were fasted the night prior to the with liberal access to water. Animals underwent PET imaging using ^18^F-FDG (7.91 ± 0.63 MBq) administered intravenously via the tail vein. Fifteen minutes before injection of ^18^F-FDG, the animals were injected intraperitoneally with 500 mg of ACLYi or an equal volume of vehicle. A 15-min PET acquisition was performed using the nanoScan PET/CT (Mediso) 45 min after tracer injection. PET images were acquired within one field of view. CT was acquired immediately following the PET for anatomical co-registration. CT acquisition parameters included 480 projections, 50-kVp tube voltage, 600 mA, 300-ms exposure time, 1:4 binning, and helical acquisition. Cardiac apical FDG signal was normalized to the blood FDG signal.

### *In Vivo* Mouse Models

C57BL/6J mice (Jackson Labs, #000664) were housed in the JHU Animal Care facilities at 22°C room temperature in 45% humidity under 12 hour light/dark cycles. Unless specified otherwise, mice were fed a standard Chow diet and had access to normal tap water *ad libidum*.

All studies were performed in compliance with the relevant ethical regulations for animal testing and research and all protocols were approved by the Johns Hopkins School of Medicine Committee on Animal Care (IACUC).

*HFpEF model:* C57BL6J mice were fed a commercially-available high-fat diet, containing 60% kilo calorie content from fat (Research Diets #D12492) and water supplemented with 0.5g/L of Nω-nitro-L-arginine methyl-ester (L-NAME, Chem-Impex #03026), as previously described^38^.

*Pressure-overload model*: mice were subjected to transaortic constriction using a 27G needle at 8-10 weeks of age, using a previously described protocol ^39^. Sham littermate controls underwent similar surgery without aortic ligature constriction.

*Tamoxifen-inducible cardiomyocyte-specific ACLY knock-down*: *Acly^Flx/Flx^* mice (Jackson Labs, MMRC stock #43555) were bred with αMHC-MerCreMer (Jackson Labs, #005657). Cardiomyocyte-specific ACLY knockdown was induced by providing 10-week-old mice with 80 mg/kg tamoxifen diet (Inotiv #TD.130858) for 8 days, after which they returned to regular diet. Age-matched αMHC-MerCreMer mice without floxed *Acly* served as controls. Echocardiography was performed 8 weeks after completion of the tamoxifen protocol.

*AAV9-ACLY shRNA induced ACLY knock down model*: C57BL6J mice aged 10 +/-2 weeks received retro-orbital injection of AAV9 expressing mouse *Acly-*shRNA or scramble shRNA. Ready-to-use AAV9 vectors were purchased from Vector Biolabs. Each mouse received a dose of 2 x 10^11^ gc of AAV9. Mice were maintained in routine housing and feeding conditions. Four weeks later, subgroups of mice were randomly assigned to b-nicotinamide mononucleotide (200 μM of NMN, Chem-Impex #33732) or vehicle added to their drinking water. Echocardiography was performed 8-10 weeks after AAV9 inoculation in all animals.

### Mouse echocardiography

M-mode images (short axis, mid-ventricular level) were obtained in conscious mice, and included LV anterior and posterior wall thickness (LVAW,d; LVPW,d), LV diameter at end-diastole and end-systole (LVIDD, LVESD), LV fractional shortening (FS) and LV ejection fraction (EF). Diastolic parameters were assessed in sedated mice using 3% inhaled isoflurane for induction followed by 1-2% titrated to maintain heart rate at 400-450 bpm. Doppler inflow velocity across the mitral valve in early (E) and late (A) diastole, peak tissue Doppler velocity at the mitral valve annulus (E’) were obtained from apical four-chamber views. Images were analyzed with VisualSonics software and average values from at least 5 heart beats per animal are reported.

### Human endomyocardial tissue

Human myocardial tissue was provided from a biobank maintained at Johns Hopkins University (HFpEF), and from the Division of Cardiology University of Pennsylvania and Gift of Life Foundation (HFrEF and non-failing donor controls) as previously described ^29^. All tissue was from the mid-septum accessed from the RV side, flash frozen at time of biopsy, and stored in - 160° C freezers. All human tissue was obtained under respective IRB approved protocols/procedures at the respective institutions. Details of the transcriptomic analysis of these cohorts has been previously reported ^29^. A summary of their clinical characteristics is provided in Supplemental Table 1. None of the patients were treated with an ACLY inhibitor (e.g. bempeproic acid). While biopsies from different patients were used for immunoblotting in the current study, they are from the same sources and reflect the same groups.

### Statistical analyses

Statistical analyses were performed in Prism (GraphPad, Ver 10) with each statistical test, median, standard deviation and number of samples per group indicated in the corresponding figure legends and panels. Normality of data was determined using the Kolmogorov-Smirnov. When normality was met, the unpaired two-tailed Student’s t test and the one or two-way Analysis of Variance (ANOVA) with Tukey’s or Sidak’s post tests were used to compare the mean values of 2 groups and 3 or more groups, respectively. For non-normally distributed data, non-parametric tests were used (Mann Whitney, or Kruskal Wallis tests). Linear regression analysis of cardiac ACLY copy number based on the patients’ pulmonary capillary wedge pressure (PCWP) was carried out in Prism using a simple linear regression model. The slopes of the regression lines were calculated and compared with 0. Statistical significance was accepted with a P value < 0.05.

## Results

### ACLY expression is depressed in human heart failure

Myocardial *ACLY* gene expression previously determined by RNAseq of human endomyocardium^29^ was significantly lower in HFrEF and HFpEF versus control; more so in HFpEF (**Fig 1A**). ACLY protein expression was reduced by 40-50% less in the HF groups (**Fig 1B**). The lower *ACLY* expression in HFpEF versus HFrEF was independent of pulmonary capillary wedge pressure at the time of biopsy, a reflector of cardiac congestion (P=1.3e-14 for between group offset, **Fig. 1C**). Using multiple regression, associations of *ACLY* (Log2-normalized reads) with age, sex, body mass index (BMI), left ventricular (LV) mass index, LV ejection fraction, estimated glomerular filtration rate (GFR), history of diabetes mellitus (DM), and history of coronary artery disease were assessed. BMI was most negatively correlated (P=2e^-6^), with borderline significant influence from GFR (P=0.05) and DM history (0.06). BMI was significantly greater in the HFpEF cohort (**Fig 1D**). *Acly* transcription is primarily regulated by carbohydrate response element binding protein (CHREBP) ^40^ and sterol regulatory element-binding protein-1 (SREBP1)^41, 42^. Gene expression of both were lower in HFrEF and HFpEF compared to controls. *SREBP2* expression was slightly higher in HFrEF but not different from controls in HFpEF (Fig. 1E). These data show ACLY expression is reduced in HF and more so in HFpEF with marked obesity.

**Figure 1:**
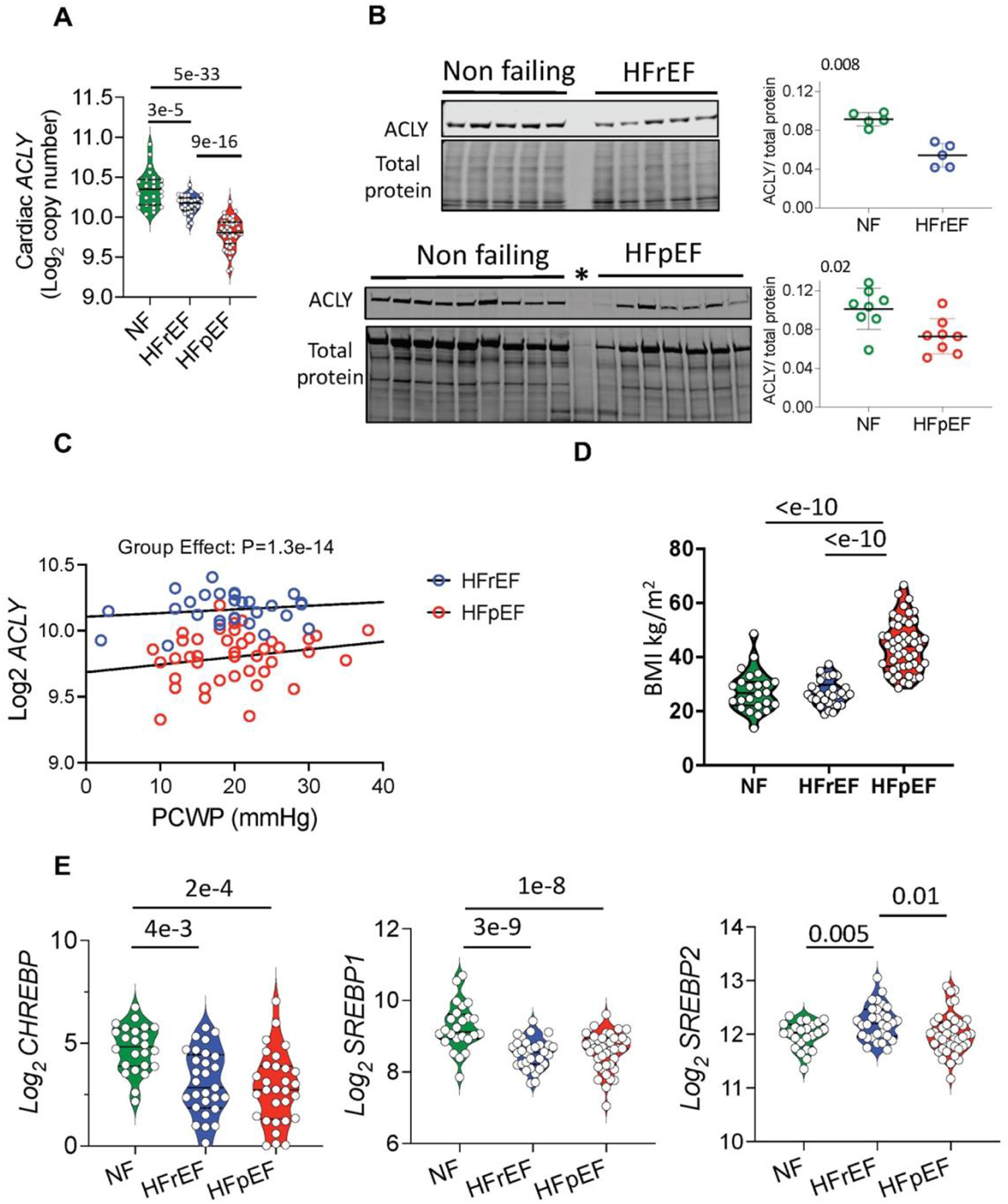
ATP citrate lyase (ACLY) is depressed in human heart failure. **A**: Cardiac mRNA levels of *ACLY* in endomyocardial biopsies from non-failing and failing human hearts (with reduced or preserved ejection fraction, HFrEF, HFpEF) from a prior study^29^. Log_2_ normalized read counts are shown based on differential expression analysis (DESeq2), and adjusted P values (Benjamini-Hochberg) reported. N= 24 NF, 30 HFrEF and 41 HFpEF patients. **B**: Immunoblot blot showing ACLY protein levels in NF versus HFrEF and NF versus HFpEF myocardium. Data are normalized to total protein and compared by Mann-Whitney test. N= 5 NF versus 5 HFrEF and N= 8 NF versus 8 HFpEF. * identifies a sample in the gel with negligible protein loading, excluded from analysis. **C:** Log_2_ *ACLY* expression versus pulmonary capillary wedge pressure (PCWP) in HFrEF (blue) and HFpEF (red) myocardium. Analysis of covariance identified a group impact but no correlation with PCWP. **D:** Comparison of body mass index (BMI) in the three cohorts, analysis by 1-way ANOVA. **E:** Cardiac mRNA levels for carbohydrate response element-binding protein (*CHREBP*) and the sterol regulatory element binding proteins 1 and 2 (*SREBP1* and *SREBP2*). Differential expression analysis, adjusted p values, and sample sizes are as determined in panel A.

### Stress regulators of myocyte/heart ACLY gene and protein expression

To test whether cardiac hemodynamic and hormonal stress alters ACLY expression, mRNA and protein levels were measured in ventricular myocardium from mice subjected to 6-weeks of pressure-overload. Both *Acly* mRNA and myocyte protein expression increased (**Fig 2A, 2B).** Immunoblots for ACLY yielded two bands in myocytes and myocardium, the upper one being specific based on data with gene silencing. HFpEF, which had the lowest *ACLY* expression generally combines hemodynamic stress with obesity. Both are included in a mouse HFpEF model (high fat diet + nitric oxide synthase inhibition^38^), and here ACLY expression was similar to controls (**Fig 2C**). This suggested obesity and hemodynamic/hormonal stress may alter *Acly* in opposite directions. To test this, cardiomyocytes were exposed for 24-hrs to either endothelin-1 (ET-1) or complete-lipid (CL). ET-1 increased *Acly* expression (**Fig 2D, E**), whereas CL reduced it (**Fig 2F, G**). Importantly, when myocytes were cultured with both, ET1-induced elevation of *Acly* was significantly blunted (**Fig 2H**). ACLY protein expression also declined in cardiomyocytes exposed to CL, and this was associated with depressed *Acly* promoter activity and reduced expression of the *Srebp1* (**Fig 2J**). Together, these findings show opposing effects of pathological hormonal or hemodynamic stress versus lipid stress on *Acly* expression, the former augmenting the latter reducing expression.

**Figure 2:**
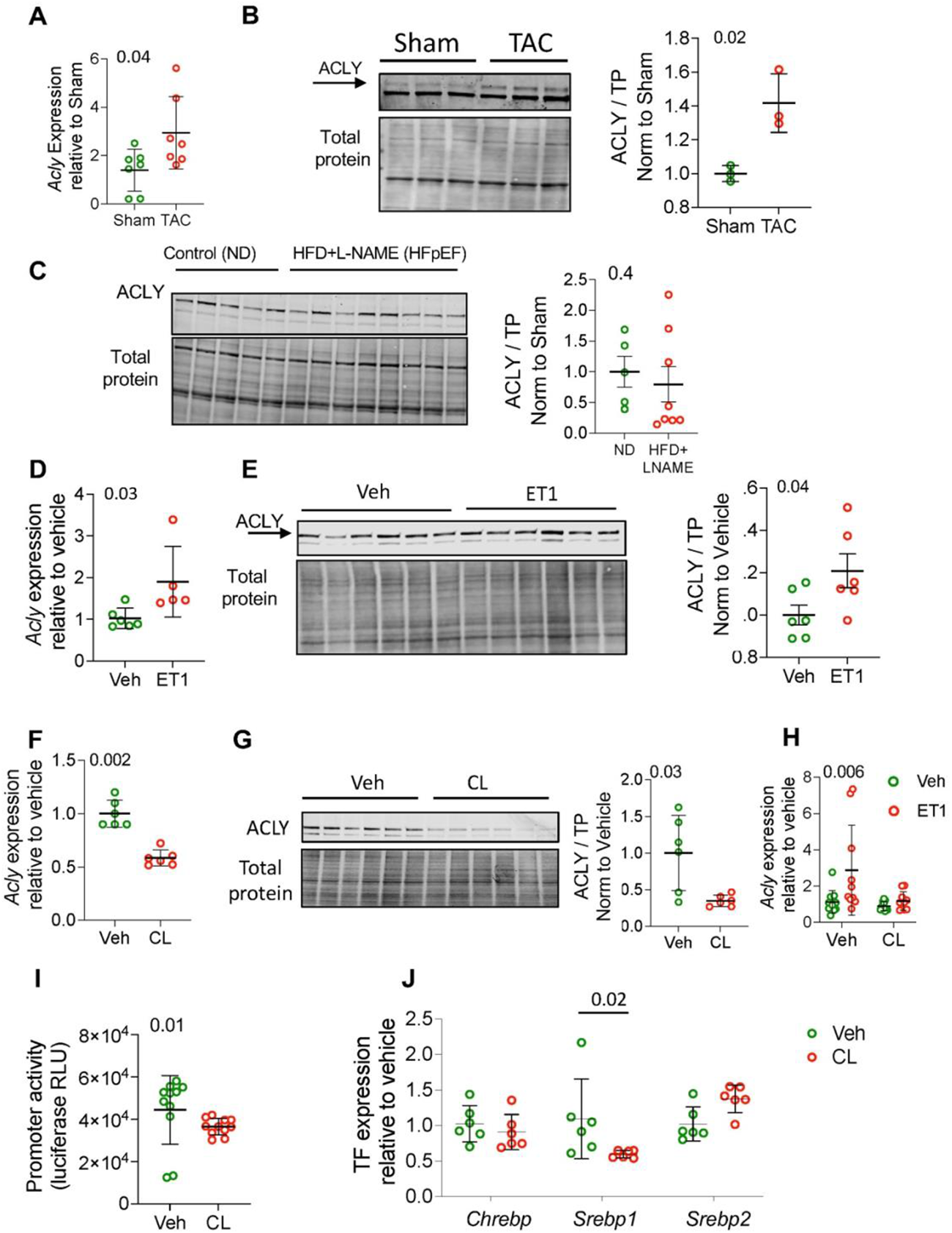
ACLY is transcriptionally upregulated in response to pro-hypertrophic stress but depressed in lipid-rich conditions. **A:** Relative change in *Acly* mRNA in myocardium from control and pressure-overload (6-week trans-aortic constriction, TAC) mice. Mann-Whitney test. N= 7 mice per group. Gene expression normalized to *Rps18* for this and all other qPCR analyses shown. **B:** Imunoblot of ACLY protein expression in cardiomyocytes isolated from control and TAC mice, and summary densitometry (Mann-Whitney test, n= 3/group). Protein expression normalized to total protein (TP) for this and all other immunoblots shown. The upper band (arrow) is ACLY. **C-**Immunoblot and summary densitometry for ACLY protein in myocardium from mice fed normal diet (ND, n=5) versus high fat diet + L-NAME (HFD+LNAME, n=8) a model of HFpEF; Mann-Whitney test. **D:** *Acly* gene and **E:** ACLY protein normalized to vehicle control in NRCMs treated with vehicle (Veh) or endothelin 1 (ET1) for 24 hours. Mann-Whitney test, N=5-6 per group. **F:** Change in *Acly* gene and **G:** ACLY protein in NRCM exposed to complete lipid (CL) for 24 hours versus vehicle control. Mann-Whitney test. N=6 per group. **H:** Relative change in *Acly* in NRCM pre-treated with vehicle or CL then exposed to ET1 or vehicle for 24 hours. 2-way ANOVA. N=10-11 per group. **I:** ACLY promoter activity by luciferase activity in NRCM in response to vehicle or CL for 2 hours. Mann-Whitney test. N=11 per group. **J:** Gene expression for transcription factors (TF) carbohydrate response element protein (*Chrebp*), sterol regulatory element binding protein 1 and 2 (*Srebp1* and *Srebp2*, respectively) in NRCM treated with vehicle or CL for 24 hours. Data normalized to vehicle control; 2-way ANOVA with Sidak’s multiple comparisons test. N=6 per group. Data displayed are all mean ±SD.

### ACLY inhibition acutely depresses cardiomyocyte and intact heart function

To test effects of acute ACLY inhibition, the selective antagonist BMS303141 (ACLYi) was applied to freshly isolated AMCMs at 1-50 μM, the high doses being well-tolerated in various proliferating cell types ^19, 43^. Nearly 80% of myocytes died after 1 hr of 50 μM ACLYi, and while viable cells persisted at 10 μM, they were often contracted as occurs before death (**Fig 3A**). Dose dependent effects of ACLYi for 1 or 24 hours were further assayed in AMCMs and NRCMs by LDH release to assess cytoxtoxicity. LDH rose significantly by one hour of ≥10 µM in AMCM and ≥ 25 µM in NRCM. (**Fig 3B**), and this toxicity worsened at 24 hours in both groups (**Fig S1a**). In contrast, ACLYi was not toxic to cardiac fibroblasts at similar exposures (**Fig S1b**).

**Figure 3:**
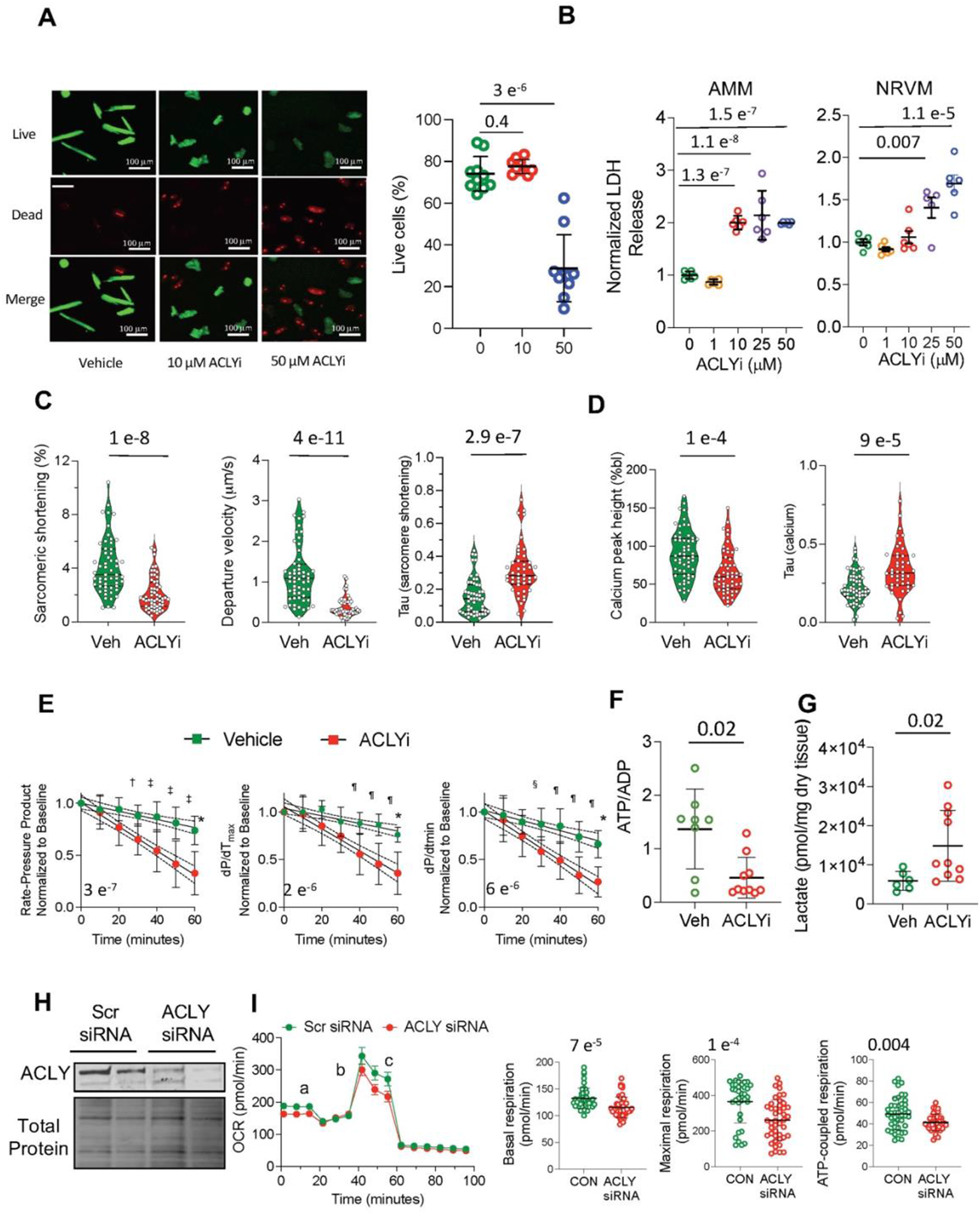
Acute ACLY inhibition is cardiotoxic and leads to cardiac dysfunction. **A:** Representative images of AMCM stained with calcein (green; live cells) and ethidium homodimer-1 (red; dead cells) following 1 hour incubation with vehicle, 10 μM or 50 μM BMS303141 (ACLYi). Percent live cells are shown at each ACLYi dose, Kruskal-Wallis test, N= 10 fields per group. **B:** Relative change in lactate dehydrogenase (LDH) released by AMCM and NRCM exposed to vehicle or ACLYi at various concentrations for 1 hour. Comparisons by one-way ANOVA with Dunnett’s multiple comparisons test. **C:** Sarcomere shortening, shortening (departure) velocity, re-lengthening time constant (tau, sarcomere shortening) and **D:** Calcium peak transient height and calcium reuptake time constant (tau, calcium) in AMCM following 1-hr treatment with vehicle or 1 μM of ACLYi. Analysis by students T-test, N=55 control, 57=ACLYi. **E:** Rate-pressure product, and maximal (dP/dt_max_) and minimal (dP/dt_min_) rate of pressure change in beating isovolumic *ex vivo* hearts exposed to vehicle or 1 μM ACLYi. Analysis of covariance; P-value for difference in slopes (lower left), and differences at individual time points determined by 2-way ANOVA with Sidak’s multiple comparisons test. N=8-9 hearts per condition. **F:** ATP/ADP ratio measured in same study after 1 hour vehicle (n=8) vs 1 μM ACLYi (n=10); Mann-Whitney test. **G:** Lactate in same study, n=6 vehicle, n=10 ACLYi, Mann-Whitney test.. **H:** Example immunoblot showing knockdown of ACLY by siRNA. **I:** Oxygen consumption rate (OCR) in NRCM transfected with scrambled or siRNA to *Acly.* Data obtained at baseline and after injections of oligomycin (a), FCCP (b) and rotenone + antimycin (c). Basal respiration, maximal respiration and ATP-coupled respiration are compared by Mann Whitney test; n= 46 per condition. All data shown as mean ±SD.

ACLYi also acutely impaired myocyte and whole heart function. AMCMs were exposed to 1 µM ACLYi for 1 hour, then electrically stimulated at 1 Hz, and sarcomere shortening and calcium transients determined. With ACLYi, the magnitude and rate of sarcomere shortening declined while cell re-lengthening (relaxation) time constant increased (**Fig. 3C**), while calcium transients had a lower peak amplitude and slower decay (**Fig 3D**). Analogous acute functional depression was observed ex vivo in mouse hearts contracting iso-volumetrically at 330 bpm. ACLYi (1 µM) induced a gradual decline in systolic (rate-pressure product and isovolumic dP/dt_max_) and diastolic (dP/dt_min_) function over time (**Fig. 3E**). With 10 µM ACLYi, developed pressure virtually ceased by 10 minutes (data not shown). These results show that myocytes and hearts require ACLY activity for normal contractile function.

At the one-hour time point, *ex vivo* volume-loaded hearts had reduced myocardial ATP/ADP ratio (**Fig. 3F**) and elevated lactate (**Fig. 3G**), suggesting depressed mitochondrial function. Myocardium was therefore assayed by targeted metabolomics, finding a significant decline in citrate with ACLYi (**Fig S1C**), but little change in other TCA intermediates. We tested whether bypassing citrate by co-administering mitochondrial permeable dimethyl-α−ketoglutarate (DMKG) would enhance downstream intermediates and restore function despite ACLYi. DMKG augmented TCA intermediates downstream of citrate substantially, but only modestly increased citrate (**Fig S2A**) and did not rescue cardiac depression by ACLYi (**Fig S2B**). The modest change in citrate despite substantial rise in other TCA intermediates suggests some functional impairment of citrate synthase with ACLYi. However, this was not an off-target effect by the inhibitor itself, as this had no impact on recombinant citrate synthase activity at 1 or even 10 µM (**Fig S2C)**.

To test if ACLYi reduced mitochondrial respiration, NRCMs were incubated with either *Acly*-siRNA or scrambled vector for 48 hours and respiration assessed (**Fig 3H**). Both basal and peak O_2_-consumption and ATP-coupled respiration were significantly reduced in si*Acly* treated cells, supporting less oxidative metabolism (**Fig. 3I**). This was not due to electron transport chain uncoupling, confirmed by unaltered TMRM fluorescence (**Fig S3A**), nor a change in glycolysis as indicated by similar extracellular acidification rates in the michondrial assay (**Figure S3B**).

### Myocyte ACLY integrates with the TCA cycle and controls NAD+/NADH redox balance

To determine whether in cardiomyocytes, cytosolic citrate metabolized by ACLY is in turn returned into mitochondria to bypass TCA cycle steps, AMCMs were incubated with uniformly labeled (M6) ^13^C glucose and then exposed to ACLYi or vehicle. As shown in (**Fig 4A**), M6-glucose is converted by glycolysis into two M3-pyruvates, which can be reduced to M3-lactate or decarboxylated in mitochondria to form M2-acetyl-CoA. M2-acetyl-CoA in turn enters the TCA cycle to generate M2-citrate and ultimately M2-malate. M2-Citrate that exits mitochondria is converted by ACLY into M2-acetyl-CoA, leaving unlabeled oxaloacetate. This is then converted to unlabeled malate (M0-malate), which can re-enter mitochondria by the malate/aspartate shuttle. Thus, if an ACLY-dependent cycle is present, the M2 malate/M2 citrate and M2 malate/M0 malate ratio should increase after ACLYi; this was indeed observed (**Fig 4B**). TCA cycle iso-topologue fractions for TCA intermediates are shown in **Fig S3B**. Notably M3-pyruvate and M3-lactate fractions were unchanged by ACLYi, further supporting no change in upstream glycolysis. Glucose uptake by the heart was assessed by *in vivo* FDG-PET imaging in mice treated with ACLYi or vehicle and was similar between groups (**Fig. S3C**). These findings support an ACLY-dependent cytosolic-to-mitochondrial cycle in myocytes that integrates with the TCA cycle.

**Figure 4:**
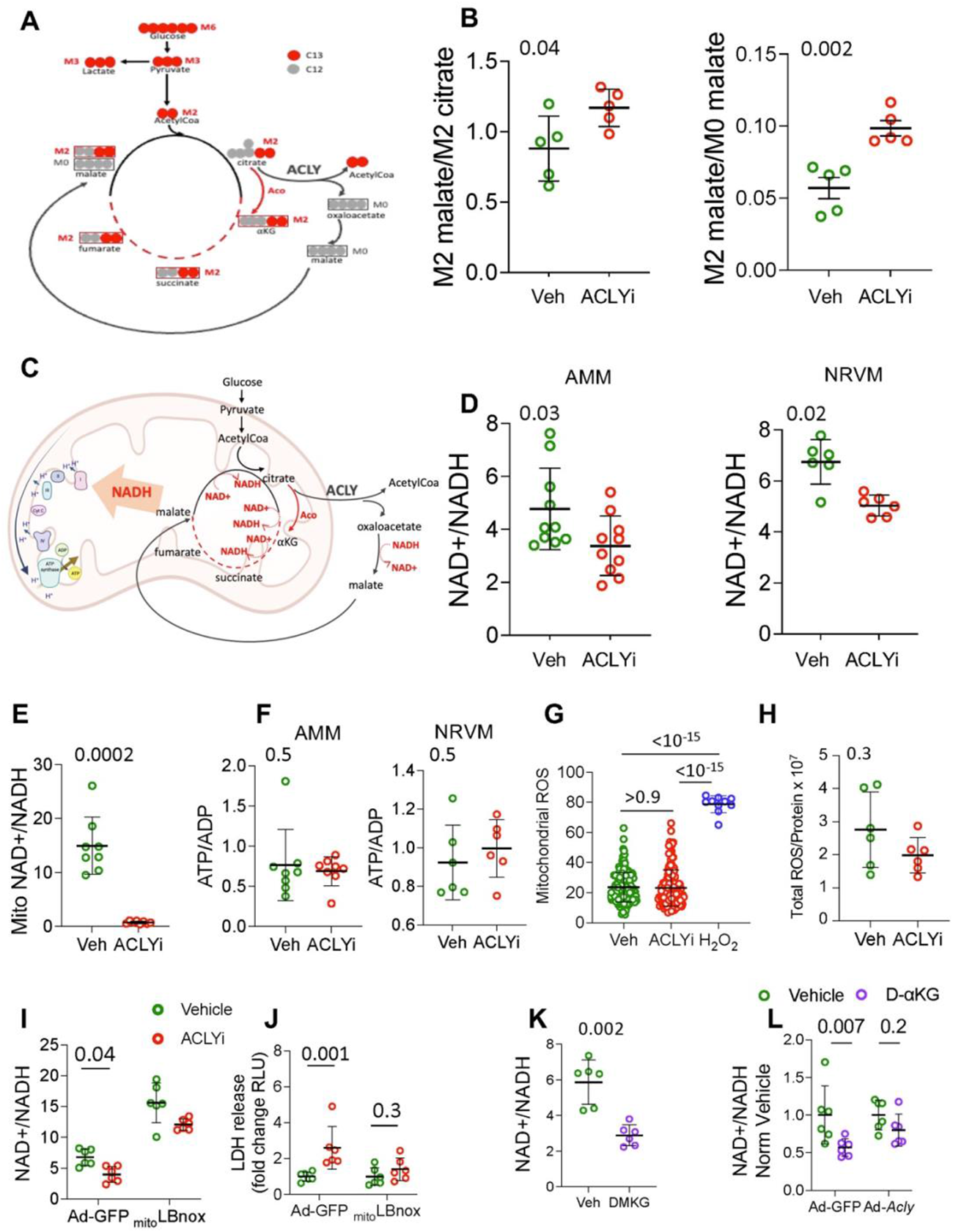
The TCA cycle pool partially flows through ACLY in cardiomyocytes to maintain the mitochondrial NAD/NADH balance. **A:** Schematic of the metabolism of fully-labeled (M6) ^13^C-glucose (U-C13 glucose) upon entering the TCA cycle. U-C13 glucose generates M2 acetylCoA, which yields M2 TCA cycle intermediates (citrate, KG, succinate, fumarate and malate) if processed through aconitase. If processed via ACLY, this results in M0-labeled malate. Upon re-uptake into mitochondrial, this reduces the net labeled levels of malate. The greater the M2 malate/M2 citrate and M2 malate/M0 malate ratio indicates more activity of intrinsic (canonical) TCA cycle via aconitase. **B:** M2 malate/M2 citrate and M2 malate/M0 malate in cardiomyocytes with or without 10 μM ACLYi for 1 hour. Analysis by Mann Whitney test. N=5 biological replicates per group. **C:** Schematic showing NAD+ and NADH species generated by the TCA cycle and anticipated changes in NADH species resulting from the activity of ACLY. Steps bypassed by the activity of ACLY are generation of 2 molecules of NADH in mitochondrion and consumption of one NADH in the cytosol. **D:** NAD+/NADH ratio in adult mouse cardiomyocytes (AMCM) and neonatal rat ventricular cardiomyocytes (NRCM) after 1 hour treatment with vehicle or 10 µM or 25 µM ACLYi respectively. Analysis by Mann-Whitney test. N=5-10 biological replicates per group. **E:** NAD+/NADH ratio in the mitochondrial compartment from NRCM after 1 hour of incubation with 25 μM ACLYi. Mann-Whitney test. N=8 biological replicates per group. **F:** ATP/ADP ratio in AMCM and NRCM after 1 hour of ACLYi. Mann-Whitney test. N=6-8 biological replicates per group. **G:** Mitochondrial ROS assessed by MitoSOX intensity in NRCM after 1 hour of incubation with vehicle, 25 μM ACLYi, or hydrogen peroxide (H_2_O_2_). Welch’s ANOVA test with Dunnet’s multiple comparisons. N=160 cells/condition and n=10 for H_2_O_2_. **H:** Cellular ROS assessed by electron paramagnetic resonance, data normalized to protein concentration in NRCM treated with vehicle or 25 μM ACLYi for 1 hour. Mann-Whitney test. N=6 biological replicates per group. **I:** Total cellular NAD+/NADH ratio and **J:** Lactate dehydrogenase release in NRCM transduced with GFP-expressing adenovirus (CON) or _mito_LBnox-expressing adenovirus after 1 hour of treatment with vehicle or 25 μM ACLYi. N=6 biological replicates/condition. 2-way ANOVA, Sidaks multiple comparisons test. **K:** NAD+/NADH in NRVM cultured in medium supplemented with 2.5 mM of dimethyl-α-ketoglutarate (DMKG) or vehicle. Mann-Whitney test. N=6 biological replicates per condition. **L:** NAD+/NADH in cells over-expressing ACLY or control (GFP) adenovirus, and exposed to vehicle or DMKG. 2-way ANOVA. N= 6 biological replicates per group. All data shown mean ±SD.

The TCA cycle generates 3 NADH while ACLY consumes 1 NADH (**Fig 4C**). Thus, the ACLY-dependent citrate-to-malate bypass generates two fewer NADH in the mitochondria versus the canonical TCA, and consumes one cytosolic NADH molecule. ACLYi should therefore increase NADH and thus lower the NAD+/NADH ratio. This prediction was confirmed after only 1 hour of ACLYi in AMCMs (*left*) and NRCMs (*right*) (**Fig. 4D**), and this ratio was more profoundly reduced in isolated mitochondria from ACLYi-treated myocytes (**Fig. 4E**), indicating mitochondrial NADH generation exceeded the capacity of the electron transport chain. ATP/ADP ratio was unaltered by ACLYi (**Fig. 4F**), perhaps reflecting the fact that unlike the beating ex vivo heart, myocytes were not performing work but were essentially quiescent. As lower NAD+/NADH ratio after ACLYi was coupled to LDH release (cytotoxicity), this suggested reductive stress. A major mechanism by which reductive stress is thought to be cytotoxic is its induction of oxidative stress via electron leak at complex I in the electron transport chain^44^. Yet as assayed by the mitochondrial superoxide sensor mitoSOX (**Fig. 4G, Fig. S3A**), electron paramagnetic resonance (**Fig 4H**), or a cytosolic ROS sensor CytoORBP (**Fig. S3B**), there was no apparent rise in ROS with 1 hr ACLYi despite the observed cardiotoxicity. Also, ACLYi induced similar LDH release in cells treated with a reducing agent N-acetylcysteine (**Fig. S3C)**, further supporting that the cytotoxic and functional behavior with ACLYi was not ROS dependent.

To test whether ACLYi-induced cytotoxicity required reduced NAD+/NADH balance, NRCMs were infected with an adenoviral construct expressing a mitochondrially targeted LB-nox enzyme (_mito_LBnox), a *Lactobacillus brevis*-derived protein that generates NAD+ from NADH^45^. This increased NAD+/NADH with or without 1-hr ACLYi (**Fig. 4I**), but also suppressed cytoxicity from ACLYi (**Fig. 4J**). Using an opposite approach, myocytes were infected with adenovirus expressing *Acly* (or control vector) and then exposed to vehicle or DMKG. In non-transfected myocytes or those infected with control adenovirus, DMKG significantly reduced NAD+/NADH ratio; however, this did not occur in myocytes with *Acly* overexpression (**Fig. 4K, 4L**). These findings show that ACLYi mediates myocyte toxicity by mitochondrial NADH accumulation and relative NAD+ reduction, and this is prevented by restoring NAD+/NADH balance.

### Genetic knock-down of ACLY in cardiomyocytes depresses cardiac function *in vivo*

To test the effect of ACLY inhibition on cardiac function *in vivo*, two models were generated. First, ACLY shRNA (or scrambled control) was expressed in an AAV9 vector and injected intravenously into mice. At 10 weeks post injection, ACLY protein expression in total myocardial lysate declined by nearly half (**Fig. 5A**). There was no change in body or heart weight, although lung wet-dry weight, a measure of lung edema, increased (**Fig. 5B**). Both end-diastolic and end-systolic volume increased while fractional shortening declined (**Fig. 5C**), indicating cardiac dilation and depressed function. Cardiac metabolomics found TCA cycle intermediates downstream of citrate increased, as well as lactate and pyruvate, compatible with increased TCA cycle flux (**Fig. 5D**). NAD+/NADH ratio declined in hearts receiving *Acly-*shRNA (**Fig. 5E**). To test whether the reduced NAD+/NADH from ACLY knock-down was causally related to cardiac dysfunction, a separate cohort of mice were fed the NAD precursor (nicotinamide mononucleotide, NMN) added to their drinking water starting 4 weeks after receiving either AAV9-*acly*-shRNA or AAV9-scrambled control. After 10-weeks of NMN treatment, NAD+/NADH ratios (**Fig 5F**) and cardiac function (**Fig 5G**) were equalized between the two groups .

**Figure 5:**
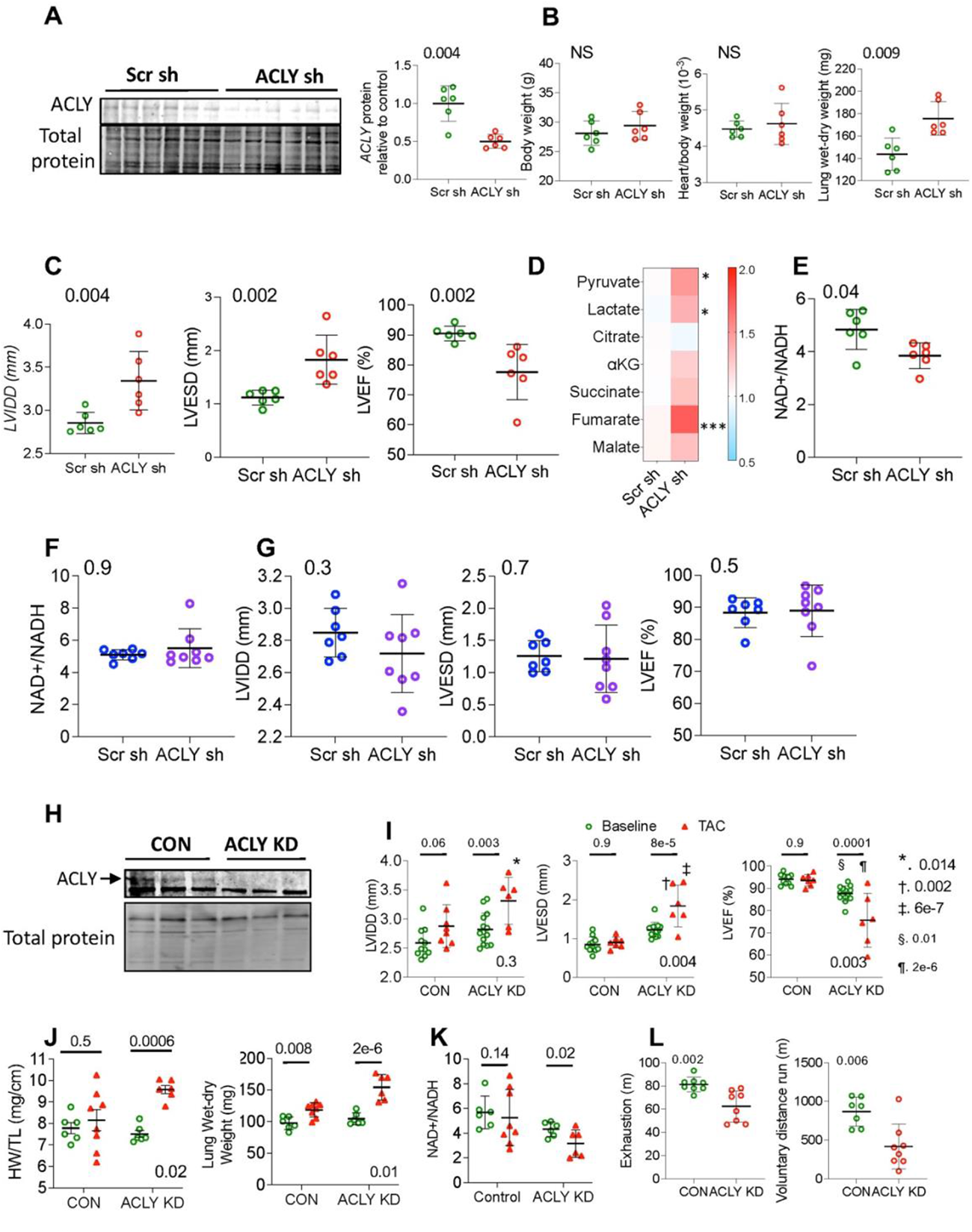
Cardiac and cardiomyocyte *Acly* gene depression *in vivo* causes ventricular dysfunction, worsened by pressure-overload, and ameliorated by NAD+ supplementation. **A:** Immunoblot blot of ACLY protein expression in myocardium from mouse model of ACLY knock down (KD) by AAV9-shRNA (ACLY sh) versus control (scramble shRNA, Scr sh). **B:** Body weight, heart weight/body weight, lung-dry lung weight and **C:** left ventricular end-diastolic and end-systolic diameters (LVIDD and LVESD) and ejection fraction (LVEF) in mice 10-weeks after receiving the respective AAV9s. **D-** Heat map for Log2-fold change in cardiac metabolites in mice 10 weeks after receiving respective AAV9s. **E-** NAD+/NADH ratio in myocardium of each respective group. Mann-Whitney test. N=6 mice/group. **F-** NAD+/NADH ratio **G-**echocardiographic parameters for AAV9 ACLY shRNA versus AAV9 scramble shRNA-treated mice concomitantly fed nucleotide mononucleotide in drinking week starting 4 after AAV9 inoculation. Mann-Whitney test. N=7-8 mice/group. **H-**Immunoblot for ACLY protein in cardiomyocytes isolated from a cardiac-specific tamoxifen-inducible ACLY Knockdown mouse model generated by breeding αMHC-MerCreMer x *Acly^Flx/Flx^* (m*Acly* KD). **I:** Echocardiographic parameters, **J-** left ventricular mass and lung wet-dry weight, and **K-** NAD+/NADH ratios between control vs m*Acly* KD groups at baseline and following 8 weeks of mild transaortic constriction (TAC). 2-way ANOVA. N=6-14 mice/group. **L-**Distance run on exhaustion test and voluntary distance run overnight are shown in mACLY KD and control (CON) mice. Mann-Whitney test. N=7-8 mice/group.

AAV9 gene delivery infects myocytes but can also impact other cell types. To further test if reduced cardiomyocyte *Acly* is sufficient to induce functional decline, a tamoxifen-inducible *Acly* knock-down model was generated, using Mer-Cre-Mer^+/-^ mice crossed with ACLY flox^+/+^ mice (m-*Acly-*KD). While total myocardial ACLY was unchanged (**Fig S5A**), isolated myocyte ACLY significantly declined (**Fig. 5H**). The m-*Acly-*KD or control mice were then subjected to mild pressure-overload by aortic constriction (mild-TAC). Significant increases in LV chamber dimensions, heart mass, and lung congestion (wet vs dry weight) and decreases in EF were only observed in m-*Acly-*KD mice exposed to mild-TAC (**Fig. 5I, 5J**). This was paralleled by depressed NAD+/NADH ratio that was also only significant after mild-TAC in the m-*Acly-*KD group. To further assess the functional impact of cardiomyocyte ACLY knock-down, mice (without TAC) were exercised by treadmill (distance traveled until exhaustion) and on a voluntary exercise wheel. Both metrics revealed a marked reduction in exercise capacity in the m-*Acly-*KD mice (**Fig. 5L**).

## Discussion

This study provides insights into the role of ACLY in the myocardium, identifying a constitutive role in supporting myocyte and intact heart function and maintaining NAD+/NADH balance. As first reported in highly proliferative cells and cells responsible for lipid biogenesis and storage^19^, ACLY also appears involved with a cytosolic circuit integrating with the TCA cycle in cardiomyocytes. This was somewhat surprising, given the primacy of the TCA cycle to cardiac energy generation, and the lack of cell proliferation or lipogenesis in cardiomyocytes. A correlate of this role is the impact of ACLY inhibition on lowering NAD+/NADH ratio, and this too was confirmed *in vitro* and *in vivo*. This imbalance was further linked to myocyte cytotoxicity and in vivo cardiac dysfunction, as both were prevented by NAD replenishment. *In vivo*, pressure-overload was associated with ACLY upregulation, suggesting an adaptive mechanism, whereas ACLY knockdown enhanced cardiac sensitivity to pressure-overload and reduced exercise capacity. Our data show HF combined with obesity, as often found in HFpEF, is associated with the lowest *ACLY* expression, suggesting this syndrome may be one where ACLY decline is particularly relevant.

Redox dyshomeostasis in the heart has been traditionally viewed from the perspective of oxidative stress; however, various cardiac stressors can also result in excess reductive capacity^44,46,44,46^. This can occur from an overabundance of antioxidants such as reduced glutathione, hyperactivation of Nrf2-Keap1(NF-E2-related factor 2-Kelch-like-associated protein-1) signaling, or increased NADH over NAD+. NADH excess in particular is thought to induce cytotoxicity by impairing^47^ endoplasmic reticular function and protein folding that requires an oxidative micro-environment, and providing excess electrons to the electron transport chain (ETC) that in turn leads to electron leakage with superoxide formation at Complex I and depression of ATP synthesis^44^. Interestingly, ACLYi did not result in increased ROS at the mitochondria or in the cytosol, and its cytotoxicity was not impeded by a reducing agent, so greater ROS does not appear involved.

An alternative to ROS is that the decline in NAD+/NADH ratio is sufficient to impede oxidative metabolism that normally oxidizes NADH. This is compatible with reduced ATP/ADP found in the working ex vivo heart. Lower NAD+/NADH is reported in humans and animal models of HF^46, 48^ and NAD supplementation benefits the heart in several relevant models ^49–51^. While causes for reduced NAD+/NADH ratio are not fully known, one mechanism is depressed mitochondrial respiration, which is also found in HF ^52^. This would sequester NAD+ as NADH, impacting enzymes that require NAD+ in mitochondria such as sirtuins 3, 4, and 5. NADH is also a potent inhibitor of citrate synthase, the rate-limiting step in the TCA cycle, and of isocitrate dehydrogenase, so its rise can impede the TCA at this position. By reducing NAD+/NADH, ACLYi creates a condition that the myocyte would normally interpret as fuel excess, and so starts shutting down NADH generating circuits (e.g. glycolysis and TCA) and ultimately the ETC. That this is occurring despite oxygen and metabolic fuel being adequate leads to depressed function and greater sensitivity to loading and hormonal stress that demand higher oxidative metabolism.

To our knowledge, this is the first study to demonstrate cardiac functional effects from acute ACLY inhibition. Given their presence within 1 hour, these changes are unlikely related to the epigenetic roles of ACLY via nuclear acetyl-CoA and associated histone acetylation to alter gene expression^18^. Longer term suppression of ACLY *in vivo* may indeed these other aspects of regulation that will need to be further dissected in future studies. While it first seemed surprising that myocytes have a substantial component of alternative TCA cycling via ACLY to regulate NAD+/NADH, in retrospect it seems reasonable given the need to finely tune this balance along with continuous high requirements for ATP. Mitochondria-deficient tumor cells display less sensitivity to ACLY depletion than those with more mitochondria^53^, and this may be mirrored by finding substantial effects from ACLYi in myocytes but no impact on fibroblasts.

As often the case, the current findings provide answers and raise new questions for further studies. Myocyte targeted gain of function models for ACLY may be useful to test if reversing its decline in HF can help offset disease. This awaits finding an animal model with downregulated ACLY as we observed in human HF and particularly HFpEF. The status of other components of the NAD+/NADH balance also remain to be investigated, for example what happens to NAD+ consuming enzymes like sirtuin deacetylases that modify mitochondrial proteins, or Poly ADP-ribose polymerases (PARPs) that regulate DNA repair, genomic stability, and cell survival. In this regard, while the KD models display cardiac dysfunction and exertional limitations, this may entail more than the energetics/metabolic consequences. Lastly, as newer ACLY inhibitors are developed that do not require hepatic activation but act directly on the enzyme, we will need to check for any cardiac signals that may arise. While attractive for lipidemic syndromes and NAFLD, the effects of ACLYi revealed here suggest some caution maybe warranted.

