## Supplementary material for "ATP Citrate Lyase Supports Cardiac Function and NAD+/NADH Balance And Is Depressed in Human Heart Failure": Acly_supplemental_BioRiv.pdf

### Supplemental data

| **Supplementatal Table 1. Clinical characteristics of Control, HFrEF, and HFpEF groups for Transcriptomic Analysis** | | | | |
|---|---|---|---|---|
| | Control (24) | HFpEF (41) | HFrEF (30) | P value |
| Age, years | 57 (52, 63) | 62 (53, 69)††† | 50 (45, 62) | 0.003 |
| Female Sex, n (%) | 10 (42%) | 24 (59%) | 10 (33%) | 0.1 |
| Race/Ethnicity | | | | <0.001 |
| African-American, n (%) | 1 (4%) | 28 (68%)***, ††† | 8 (27%)* | |
| Caucasian, n (%) | 22 (92%) | 11 (27%) | 21 (70%) | |
| Medications | | | | |
| ACEi or ARB, n (%) | 6 (25%) | 26 (63%)** | 20 (67%)** | 0.003 |
| Beta Blocker, n (%) | 6 (25%) | 22 (54%)*, ††† | 28 (93%)*** | <0.001 |
| Loop Diuretic, n (%) | 0 (0%) | 39 (95%)*** | 30 (100%)*** | <0.001 |
| Past Medical History | | | | |
| Hypertension, n (%) | 11 (46%) | 40 (98%)*** | 30 (100%)*** | <0.001 |
| Diabetes, n (%) | 3 (12%) | 26 (63%)***, † | 9 (30%) | <0.001 |
| Coronary artery disease, n (%) | 1 (4%) | 4 (10%) | 5 (17%) | 0.36 |
| BMI, $kg/m^2$ | 27 (22, 31) | 41 (36, 46)***, ††† | 26 (23, 29) | <0.001 |
| LVEF, % | 61 (60, 65) | 65 (60, 70)††† | 18 (11, 20)*** | <0.001 |
| LV End-diastolic dimension, cm | 4.0 (3.9, 4.5) | 4.6 (4.0, 5.0) *, ††† | 6.8 (6.2, 7.3)*** | <0.001 |
| Sex-adjusted LV mass/height$^{1.7}$, $g/m^{1.7}$ | 95 (86, 115) | 107 (83, 131)† | 121 (113, 150)** | 0.004 |
| eGFR, $mL/min/1.73m^2$ | 83 (56, 104) | 48 (33, 70)**, †† | 70 (56, 82) | 0.003 |
| Invasive Hemodynamics | | | | |
| RA, mmHg | N/A | 12 (8, 15)†† | 7 (5, 11) | 0.012 |
| PASP, mmHg | N/A | 45 (33, 54) | 48 (40, 56) | 0.68 |
| PAmean, mmHg | N/A | 29 (23, 35) | 29 (23, 34) | 0.55 |
| PCWP, mmHg | N/A | 20 (15, 24) | 20 (16, 24) | 0.94 |
| CI, $L/min/m^2$ | N/A | 2.52 (2.29, 3.09)††† | 2.05 (1.80, 2.30) | <0.001 |
| PVR ≥ 3wu, n (%) | N/A | 5 (12%) | 9 (30%)** | 0.006 |

Data are n (%) or median (25th-75th percentile). Fisher's exact test used for categorical variables. Kruskal-Wallis test used for continuous variables. *p<0.05 vs Control, **p≤0.01 vs Control, ***p≤0.001 vs Control. †p<0.05 vs HFrEF, ††p≤0.01 vs HFrEF, †††p≤0.001 vs HFrEF. ACEi, angiotensin converting enzyme inhibitor; ARB, angiotensin II receptor blocker; BMI, body mass index; LVEF, left ventricular ejection fraction; LV, left ventricle; eGFR, estimated glomerular filtration rate; RAP, right atrial pressure; PASP, pulmonary artery systolic pressure; PADP, pulmonary artery diastolic pressure; PAmean, mean pulmonary artery pressure; PAWP, pulmonary artery wedge pressure; CO, cardiac output; CI, cardiac index; PVR, pulmonary vascular resistance; wu, Wood units; PA, pulmonary artery; N/A, invasive hemodynamics not available for control patients. Sex-adjusted LV mass/height$^{1.7}$ was calculated by multiplying by a constant of 1.28 for women.

**Supplemental Figures:**

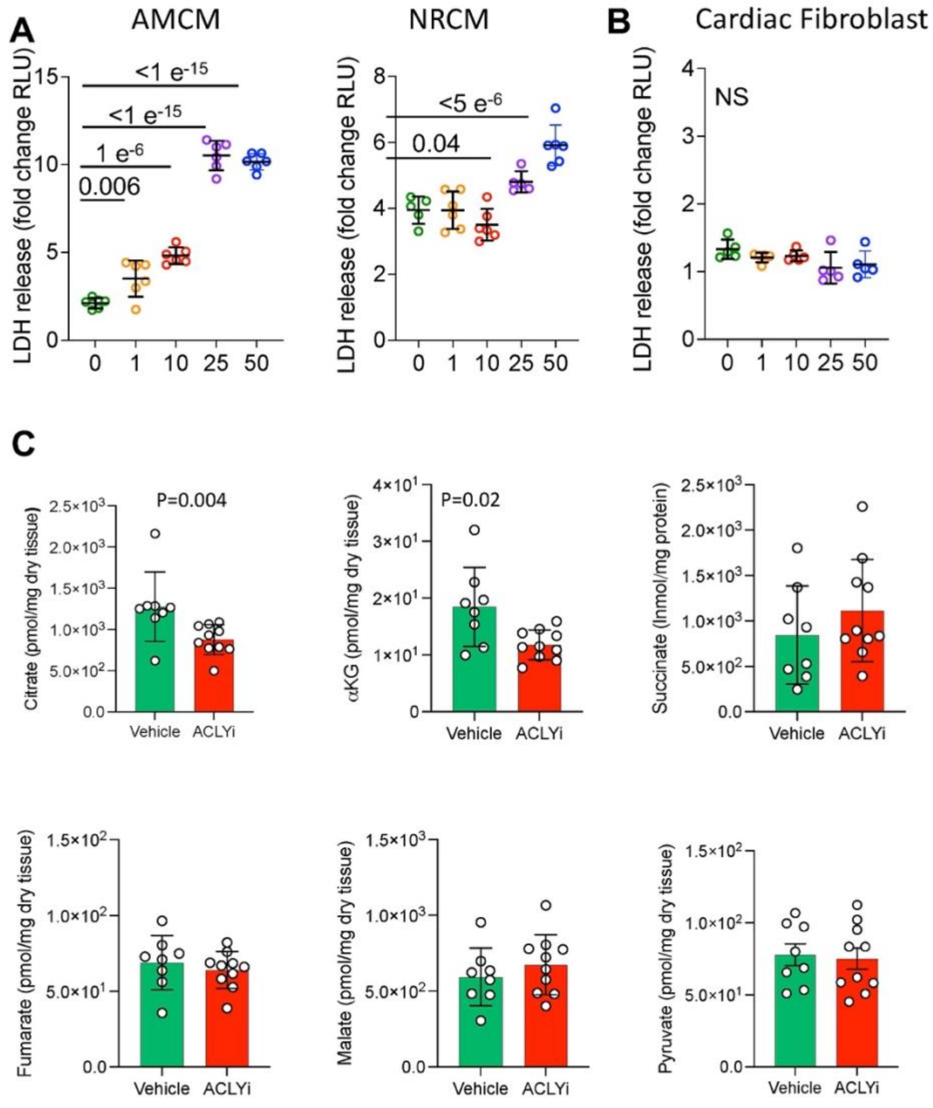


**FIGURE S1: A:** Lactate dehydrogenase (LDH) in culture medium for adult mouse cardiomyocytes (AMCM), neonatal rat cardimyocytes (NRCM), and **B:** cardiac fibroblasts exposed to either vehicle or ACLYi at varying concentrations for 24 hours. Analysis by 1-way ANOVA, p-values from Dunnett's multiple comparisons test. **C:** Concentration of TCA cycle intermediates and pyruvate level, each normalized to tissue weight from beating and loaded ex vivo hearts treated with 1 μM ACLYi versus vehicle for 1 hour. Mann-Whitney test. N=7-10 hearts/group.

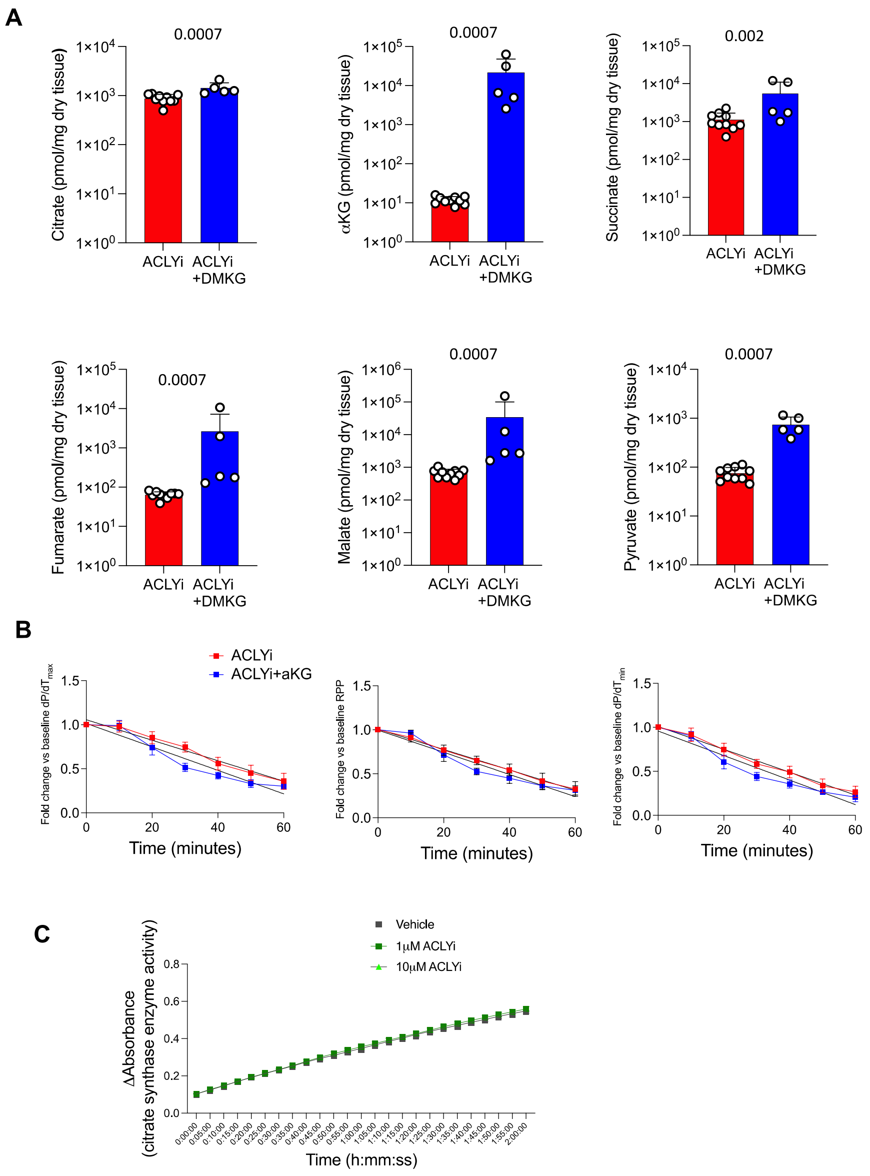


**FIGURE S2: A-** Concentration of TCA cycle intermediates and pyruvate levels, normalized to tissue weight from beating and loaded ex vivo hearts treated with 1 μM ACLYi ±dimethyl-α-ketoglutarate (DMKG) for 1 hour. Mann-Whitney test. N=10 hearts in ACLYi group and N= 5 in ACLYI +DMKG group. **B-** Change in rate-pressure product (RPP), maximal and minimal rate of pressure change ($dP/dt_{max}$, $dP/dt_{min}$) normalized to baseline for the same hearts. 2-way ANOVA with Sidak's multiple comparisons test; N=5-10 hearts per condition. **C:** Activity of recombinant citrate synthase incubated with vehicle or ACLYi at 1 μM and 10 μM. N=4 replicates/condition.

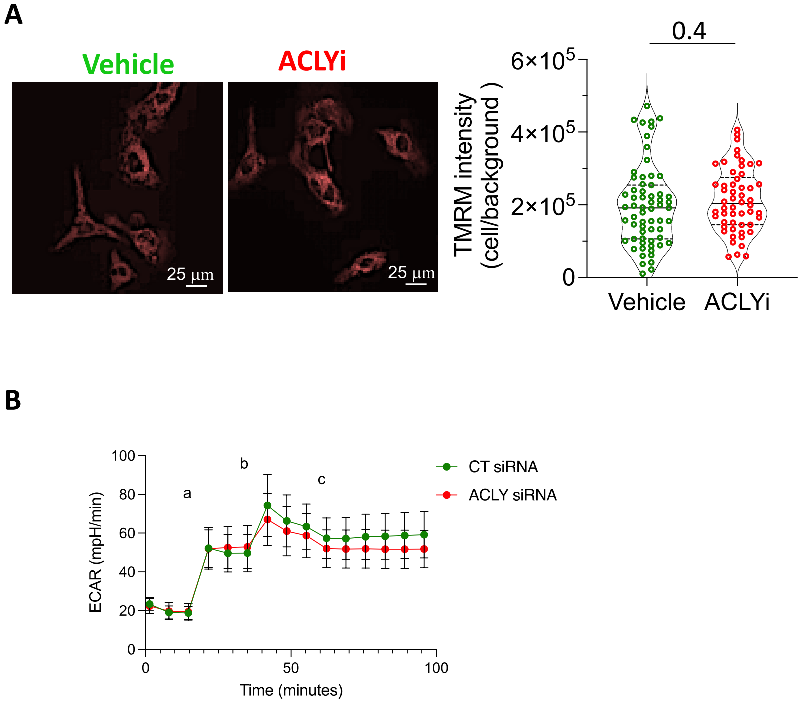


**FIGURE S3: A:** Representative confocal fluorescence microscopy images of neonatal rat cardiomyocytes (NRCM) stained with tetramethylrhodamine methylester perchlorate (TMRM) after 1 hour of treatment with 25 μM ACLYi. TMRM intensities normalized to background were compared between vehicle-treated and ACLYi-treated cells. Mann Whitney test: N= 50-60 cells/condition. **B:** Extracellular acidification rate (ECAR) from mitochondrial respiration analysis (Fig. 2I) assessing impact of *Acly* gene silencing on metabolism. ACLY inhibition reduced oxygen consumption, however as shown in this panel, there was no change in ECAR, supporting minimal impact on glycolysis. The various stages of the assay are addition of a) oligomycin (inhibit ATP synthase), b) FCCP (uncouple electron transport from ATP synthesis), and c) rotenone and antimycin A (inhibit electron transport chain).

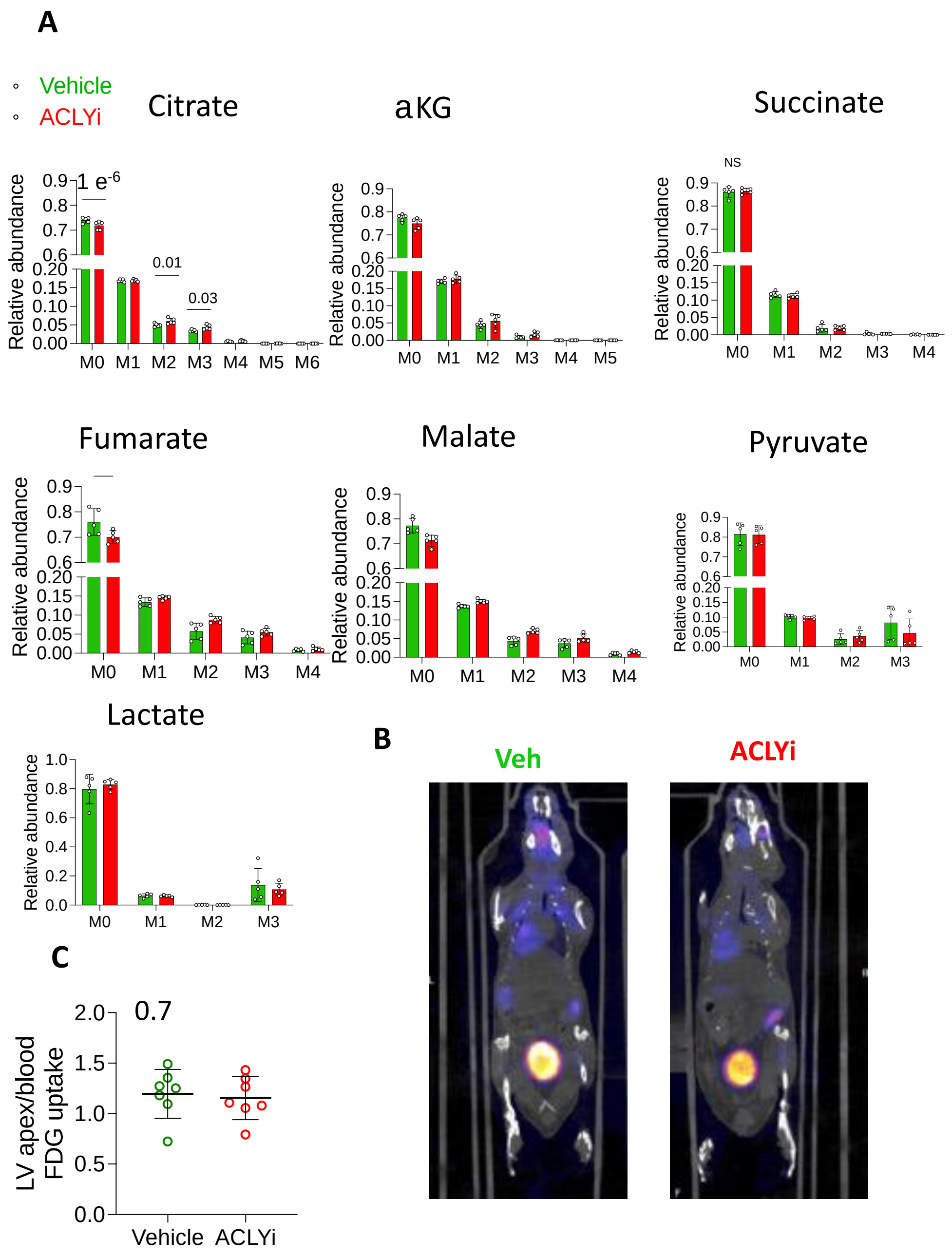


**FIGURE S4: A:** Isotopologue distribution in adult mouse cardiomyocytes (AMCM) after 1 hour of uniformly-labeled C13 glucose tracing with vehicle versus 10 µM ACLYi treatment. 2-way ANOVA with Sidak's multiple comparisons test. Only P values <0.05 are shown. **B:** Cardiac $^{18}$F-FDG uptake after 1 hour of intraperitoneal treatment with 500 µg of ACLYi or vehicle in WT C57Bl6J mice following an overnight fast. **C:** Left ventricular apex FDG uptake, normalized to blood uptake was compared using t-test. N= 7 animals/condition.

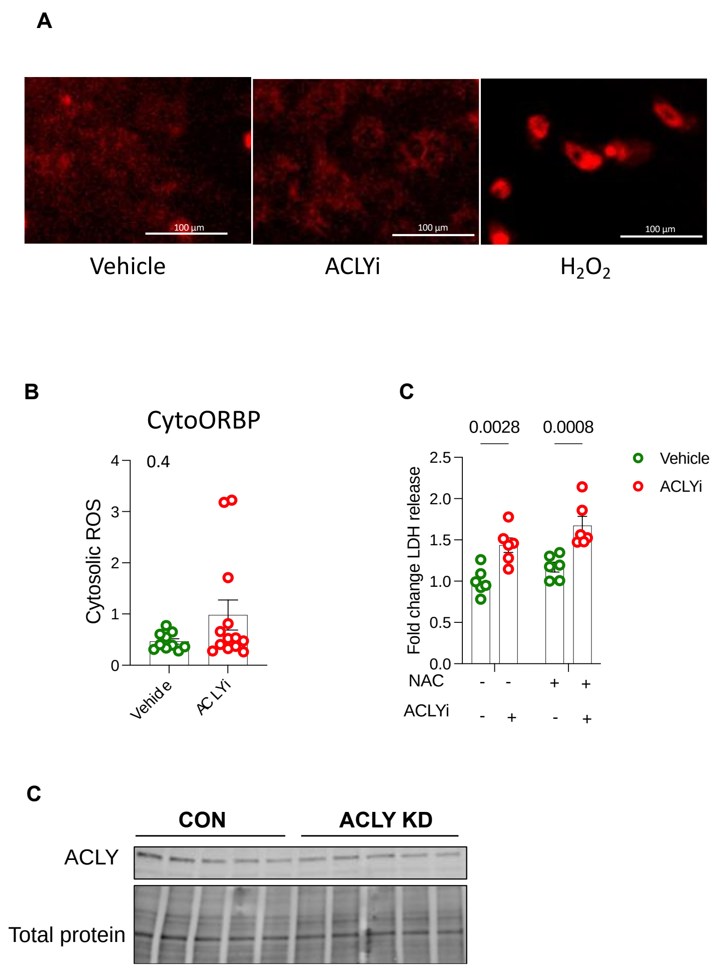


**FIGURE S5: : A-**Representative confocal microscopy images of neonatal rat ventricular myocytes (NRCM) stained with mitoSox$^{TM}$ after 1 hour of incubation with vehicle, 25 µM ACLYi or hydrogen peroxide ($H_2O_2$, positive control). **B-** Cytosolic reactive oxygen species assessed using a cytosolic ROS sensor (cytoORBP) in NRCM after 1 hour of treatment with vehicle versus 25 µM ACLYi. Comparison using the Mann-Whitney test. N=10-13 fields per condition. **C-** Relative change in the release of lactate dehydrogenase (LDH) in the culture medium by NRCM exposed to vehicle or 25 µM ACLYi for 1 hour in the absence or presence of the anti-oxidant N-acetylcysteine. **D-** Immunoblot for ACLY and total protein level from whole myocardium in αMHC-MerCreMer x *Acly*$^{Flx/Flx}$ (ACLY KD) mice versus control MerCreMer x *Acly*$^{Flx\text{-}/Fl\text{-}}$ (n=5 mice/group).
